# Protein Coronas on Functionalized Nanoparticles Enable Quantitative and Precise Large-Scale Deep Plasma Proteomics

**DOI:** 10.1101/2023.08.28.555225

**Authors:** Ting Huang, Jian Wang, Alexey Stukalov, Margaret K. R. Donovan, Shadi Ferdosi, Lucy Williamson, Seth Just, Gabriel Castro, Lee S. Cantrell, Eltaher Elgierari, Ryan W. Benz, Yingxiang Huang, Khatereh Motamedchaboki, Amirmansoor Hakimi, Tabiwang Arrey, Eugen Damoc, Simion Kreimer, Omid C. Farokhzad, Serafim Batzoglou, Asim Siddiqui, Jennifer E. Van Eyk, Daniel Hornburg

## Abstract

**Background:** The wide dynamic range of circulating proteins coupled with the diversity of proteoforms present in plasma has historically impeded comprehensive and quantitative characterization of the plasma proteome at scale. Automated nanoparticle (NP) protein corona-based proteomics workflows can efficiently compress the dynamic range of protein abundances into a mass spectrometry (MS)-accessible detection range. This enhances the depth and scalability of quantitative MS-based methods, which can elucidate the molecular mechanisms of biological processes, discover new protein biomarkers, and improve comprehensiveness of MS-based diagnostics.

**Methods:** Investigating multi-species spike-in experiments and a cohort, we investigated fold-change accuracy, linearity, precision, and statistical power for the using the Proteograph™ Product Suite, a deep plasma proteomics workflow, in conjunction with multiple MS instruments.

**Results:** We show that NP-based workflows enable accurate identification (false discovery rate of 1%) of more than 6,000 proteins from plasma (Orbitrap Astral) and, compared to a gold standard neat plasma workflow that is limited to the detection of hundreds of plasma proteins, facilitate quantification of more proteins with accurate fold-changes, high linearity, and precision. Furthermore, we demonstrate high statistical power for the discovery of biomarkers in small- and large-scale cohorts.

**Conclusions:** The automated NP workflow enables high-throughput, deep, and quantitative plasma proteomics investigation with sufficient power to discover new biomarker signatures with a peptide level resolution.

## Introduction

Proteomes are challenging to study due to their complex biochemistry and extensive dynamic range in which protein concentrations vary greatly. For instance, in plasma, 22 blood proteins constitute about 99% of the protein mass, and the remaining 1% consists of thousands of distinct lower abundant proteins and their proteoforms (1). We recently introduced a nanoparticle (NP) protein corona-based workflow that alleviates the dynamic range challenges by compressing it to a more accessible scale at the nano-bio interface (2–4). This novel combination of protein coronas and the unbiased liquid chromatography mass spectrometry (LC-MS) readout has been demonstrated to provide an unprecedented depth at scale leading to new biological insights (2,3,5).

Protein quantification in complex samples, which is essential for capturing biology, can be challenging and has been explored in multiple studies using LC-MS workflows (6–9). Quantification performance can be evaluated at the level of precision and accuracy (**Figure 1A**). Furthermore, accuracy can be dissected into three distinct facets: absolute accuracy, which pertains to protein concentration; relative fold-change accuracy, which provides an accurate estimation of protein up- or down-regulation within a given sample; and linearity, which ensures consistent patterns of up- and down-regulation across various samples for each protein (**Figure 1B**). To discover biological signals, it is primarily important that measured fold-changes follow systematic trends. For instance, the signal should be linear on a logarithmic scale and if desired, calibration functions can be employed to enhance fold-change accuracy or to determine analyte concentrations, thereby refining the overall quantification process (**Figure 1Biv)**.

**Figure 1.**
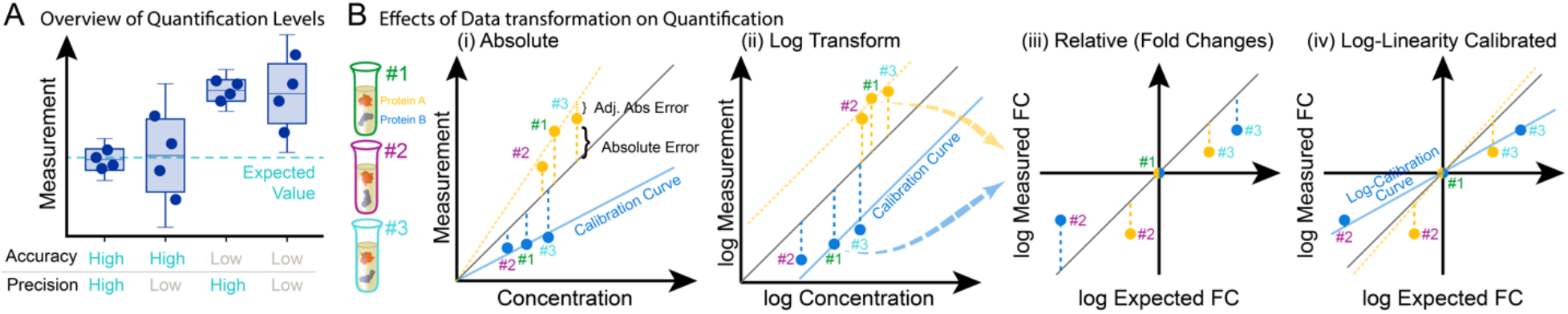
Overview of Quantification Levels and Data Transformation. (A) Two quantitative performance metrics are: 1) accuracy, measuring how close a measurement is to the true value; and 2) precision, measuring how close are the measurements across replicate analyses. (B) The measurement of two proteins (A, orange, and B, blue) across 3 biosamples (samples #1, #2, and #3) illustrates three layers of quantification accuracy: absolute accuracy (i and ii for untransformed and log-transformed data, respectively), relative fold-change (FC) accuracy (iii), and linearity (iv).

While NP-coronas can overcome the fundamental bottleneck of the plasma’s wide dynamic range by compressing it at the nano-bio interface, it is important to understand the quantification performance of this workflow. This motivated us to evaluate the proteome-wide quantification performance of the NP corona-based plasma workflow in terms of relative fold-change accuracy, linearity, and precision. We employed the Proteograph™ Product Suite with the data-independent acquisition (DIA) method to analyze a modified version of the gold standard multi-species proteome spike-in experiment (Experimental overview shown in **Figure 2A, B**) (8,10). Additionally, we investigated assay variance across a 200-sample cohort (over 1,000 LC-MS runs) to evaluate precision of the NP corona-based proteomics workflow and statistical power to detect even small fold-changes reliably.

**Figure 2.**
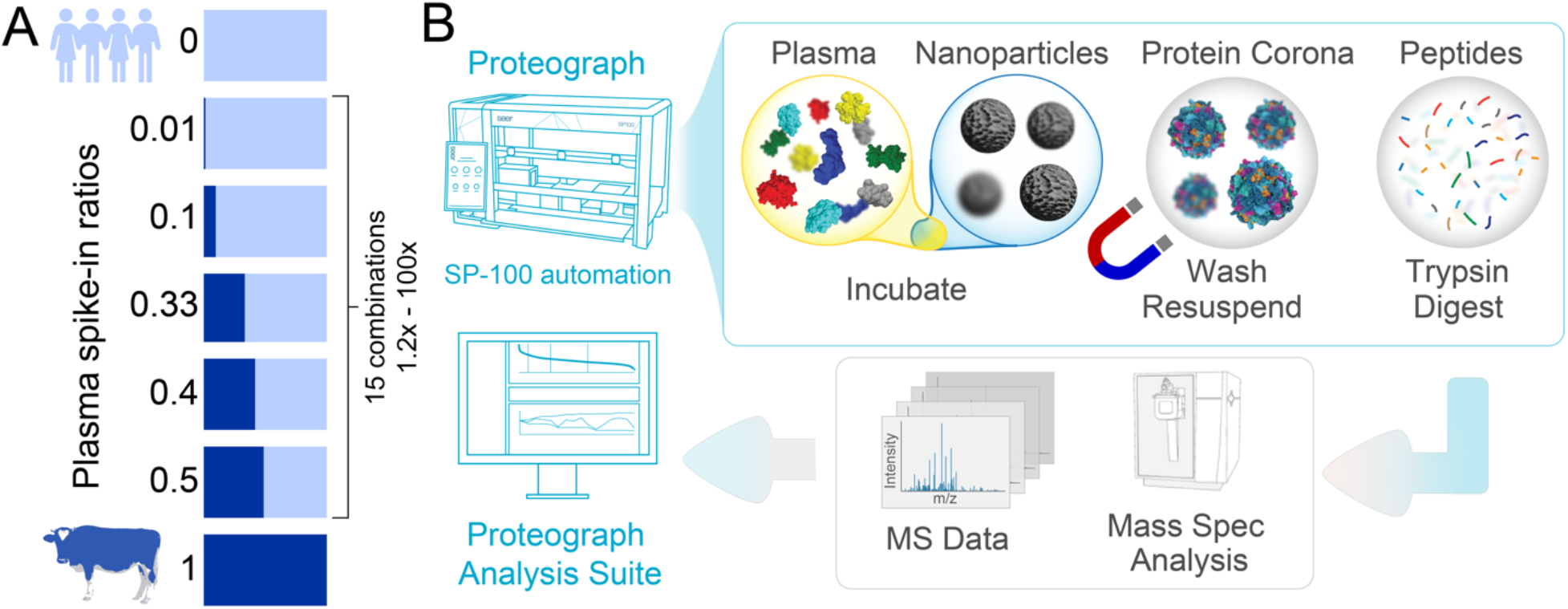
Study Overview. (A) Experimental design of the spike-in experiment, in which a bovine plasma proteome is spiked into a human plasma proteome at seven different ratios. (B) Proteograph™ workflow overview which includes protein corona formation, proteins denaturation, reduction, alkylation, protein digestion, and peptide desalting on the Proteograph SP100 automation instrument. Peptides are quantified, dried, and resuspended before injection onto an LC-MS system.

### Study design

We evaluated the quantification performance of a NP-corona based workflow by assessing relative fold-change accuracy and linearity using a modified version of the gold standard multi-proteome spike-in experiment (10,11) (**Figure 2A**). Specifically, to evaluate both large and small fold-changes, bovine plasma samples were mixed with two pooled human plasma sample (referred to as IP10 and PC6) at seven different Bovine:Human ratios (1:0, 1:1, 1:1.5, 1:2, 1:9, 1:99, 0:1). We assessed the seven Bovine:Human mixed samples using the fully automated Proteograph workflow (**Figure 2B**), as well as a reference neat plasma workflow. The neat plasma workflow provides a baseline plasma LC-MS proteomics performance since it involves minimal processing steps, no protein coronas, but is limited to high abundance proteins. Additionally, we evaluated the precision of quantification across a cohort of 200 subjects (more than 1000 MS injections) to determine the statistical power of Proteograph in cohorts.

## Materials and Methods

### Spike-in Experiment

K_2_EDTA bovine plasma (Innovative Research, USA) was spiked into two different pooled human plasma samples (IP10 collected in K_2_EDTA, and PC6 collected in anticoagulant citrate phosphate dextrose (cpd) tubes) with the following vol/vol ratios of (Bovine:Human) 1:0, 1:1, 1:1.5, 1:2, 1:9, 1:99, 0:1 in multi-day quadruplicates. Each replicate of each dilution and sample set was processed as a separate automated Proteograph™ run (V1.2, S55R1100), along with its neat plasma digest. Equal volumes of the peptide elution were dried down in a SpeedVac (3 hours-overnight), and peptides were stored at -80 °C before reconstitution in 20 ul 0.1% formic acid (FA) and 3% acetonitrile (ACN) and LC-MS analysis.

### High-Throughput Capillary Flow DIA LC-MS Analysis

For analysis on the Orbitrap Exploris 480 mass spectrometer (Thermo Fisher Scientific), peptides were loaded on an Acclaim™ PepMap™ 100 C18 (0.3 mm ID x 5 mm) trap column and then separated on a 50 cm μPAC™ HPLC column (Thermo Fisher Scientific) at a flow rate of 1 μL/minute using a gradient of 5-25% solvent B (0.1% FA, 100 % ACN) in solvent A (0.1% FA, 100% water) over 22 minutes, resulting in a 33-minute total run time. 150-400 ng of material per NP were analyzed in DIA mode using 10 m/z isolation windows from 380-1000 m/z. MS^1^ scans were acquired at 60k resolution and MS^2^ at 30k resolution.

The DIA LC-MS data were analyzed using DIA-NN v1.8 (12) in library-free mode and searched against Uniprot Human plus Bovine protein database (105,533 entries). The following parameters were used: “--mass-acc-ms1 10 --mass-acc 10 --qvalue 0.01 --matrices --missed-cleavages 2 -- met-excision --cut K*, R* --smart-profiling --relaxed-prot-inf --pg-level 1 --reannotate”; the remaining parameters were set to default. The false discovery rate (FDR) cutoffs at precursor and protein levels were set to 1%. Most analyses were based on species specific (unique) peptides and proteins. Details of downstream statistical analyses are shared in the supplementary information ‘LC-MS Data Analysis Exploris 480 MS (supplementary information)’.

### Nanoparticle Workflow LC-MS comparison

For the Orbitrap Astral mass spectrometer (Thermo Fisher Scientific), peptides were loaded onto a μPAC™ Neo trap column (Thermo Fisher Scientific) at a flow rate of 1 μL/minute using a gradient of 5-22.5% solvent B (0.1% FA, 100 % ACN) in solvent A (0.1% FA, 100% water) over 13 minutes, resulting in a 20-minute total run time. 400 ng of material per injection was analyzed in DIA using 3 m/z isolation windows from 380-980 m/z. MS1 scans were acquired at 240k resolution and MS2 at >80k resolution. For comparison search, LC-MS/MS data were analyzed using DIA-NN v1.8.1 (12) in library free mode and searched against the same Uniprot Human database (48,957 entries) as used for the Orbitrap Exploris 480 analysis (no matching between runs). The following parameters were used: “--matrices --met-excision --cut K*, R* --relaxed-prot-inf --pg-level 1 --individual-mass-acc --individual-windows “; the remaining parameters were set to default. FDR cutoffs at precursor and protein levels were set to 1%.

### Reproducibility Study

A control pooled human plasma sample was processed with two different Proteograph SP100 automation instruments, on two separate days each, resulting in a total of 4 batches (plates). Tryptic peptides from 6 replicates were analyzed using the same set up as outlined above with an Orbitrap Exploris 480 Mass Spectrometer here in DDA mode with a 30-minute LC gradient. LC-MS data files were processed using Proteograph Analysis Suite (PAS MaxQuant module), with 1% FDR at the protein and peptide levels (13). Coefficient of variance (CV) were evaluated on median normalized peptide intensities for peptides found in 3 or more replicates, with four different replicate grouping methods: within plate (batch), within day across SP100 instruments, within SP100 across days, and between days and SP100 instruments.

### Cohort Study

A pooled human plasma (process quality control (QC)) sample, consisting of pre-pooled K_2_ EDTA plasma from ProMedDx (Norton, MA) derived from healthy subjects, was processed as control on each Proteograph Assay plate analyzed across 200 plasma cohort samples (1-3 controls included on 14 plates on three Proteograph SP100 automation instruments) resulting in 117 QC measurements. Peptides were loaded on an Acclaim PepMap 100 C18 (0.3 mm ID x 5 mm) trap column and then separated on a 50 cm μPAC analytical column (Thermo Fisher Scientific) at a flow rate of 1 μL/minute using a gradient of 5 – 25% solvent B (0.1% FA, 100 % ACN) in solvent A (0.1% FA, 100% water) over 42 minutes (63-minutes total run time) on two different timsTOF Pro mass spectrometers (Bruker). A total of 200 ng of peptides per NP was analyzed in ddaPASEF mode using ion mobility range of 0.85 – 1.30 V.s/cm^2^ with 100 ms accumulation time. DDA data were analyzed with MSFragger 3.4 using the default settings in PAS, based on the Uniprot Human FASTA database (see above). Peptide intensity CVs were calculated based on median normalized intensities for peptides found in 3 or more replicates to represent the variability of performance across and within the two LC-MS systems during this cohort study. Statistical details are shared in the supplementary information ‘Cohort Study (supplementary information)’.

### Reference Spike-in Experiment

We compared the Proteograph workflow performance to published work (37), where Tognetti et al. had evaluated the performance of a plasma depletion workflow using a controlled spike-in experiment. Human plasma background was spiked with fixed ratios into Saccharomyces cerevisiae (1:1.3) and Escherichia coli (1:0.5), each in 20 technical replicates. For our comparison, we obtained the Spectronaut precursor reports for both the depletion and neat workflows from the MassIVE repository (dataset ID: MSV00088180). The precursor-level fold-changes between two conditions were calculated and compared to the spiked-in ratios of corresponding proteomes. To make the fold-change errors more comparable to the 4 replicates used in our study, we downsampled the data to four replicates randomly 100 times and took the mean fold-change error for each precursor.

## Results

### High proteome coverage and precision of NP-based workflows

Compared to the neat plasma workflow, the compression of dynamic range in NP-protein coronas results in significantly higher coverage of proteomes (2,3,14). Importantly, absolute proteome depths and fold improvements depend on plasma sample complexity and specifics of the LC-MS workflow. For instance, Proteograph in conjunction with the latest Thermo Scientific Orbitrap Astral Mass spectrometer reveals more than 6,000 proteins and 50,000 precursors in plasma compared to only hundreds of proteins accessible with a traditional upfront neat plasma workflow (**Figure 3A**). To evaluate the quantitative performance of a NP workflow, we evaluated a set of mixed species proteome experiments on the Orbitrap Exploris 480 system to a depth of more than 3000 proteins across all dilutions (**Figure 3B**). Importantly, a stringent 1% false discovery rate for the identification at precursor and protein level was applied in every analysis. Moreover, while the mixed species experiments are well established to investigate quantification, these analyses require additional filtering steps including the removal of shared proteins and peptides, or detection across the entire range of the dilutions, which reduces the reported unique protein identifications. The NP workflow provides more measurements at any quantification stringency and every spike-in ratio (**Figure 3C, D**). For example, at 1:1 spike-in ratio, a median of 268% more proteins were quantified using the NP workflow compared to neat plasma digestion (395 vs. 1,058) with the CV of protein intensities across four assay replicates below 20% (**Figure 3D**). Proteins detected in both, NP-based and neat plasma digestion workflows, show similar or slightly increased CVs on Proteograph (**Figure 3D**). Overall, these results demonstrate superior NP performance yielding more proteins than neat plasma alone with high precision and for shared proteins, showing comparable reproducibility.

**Figure 3.**
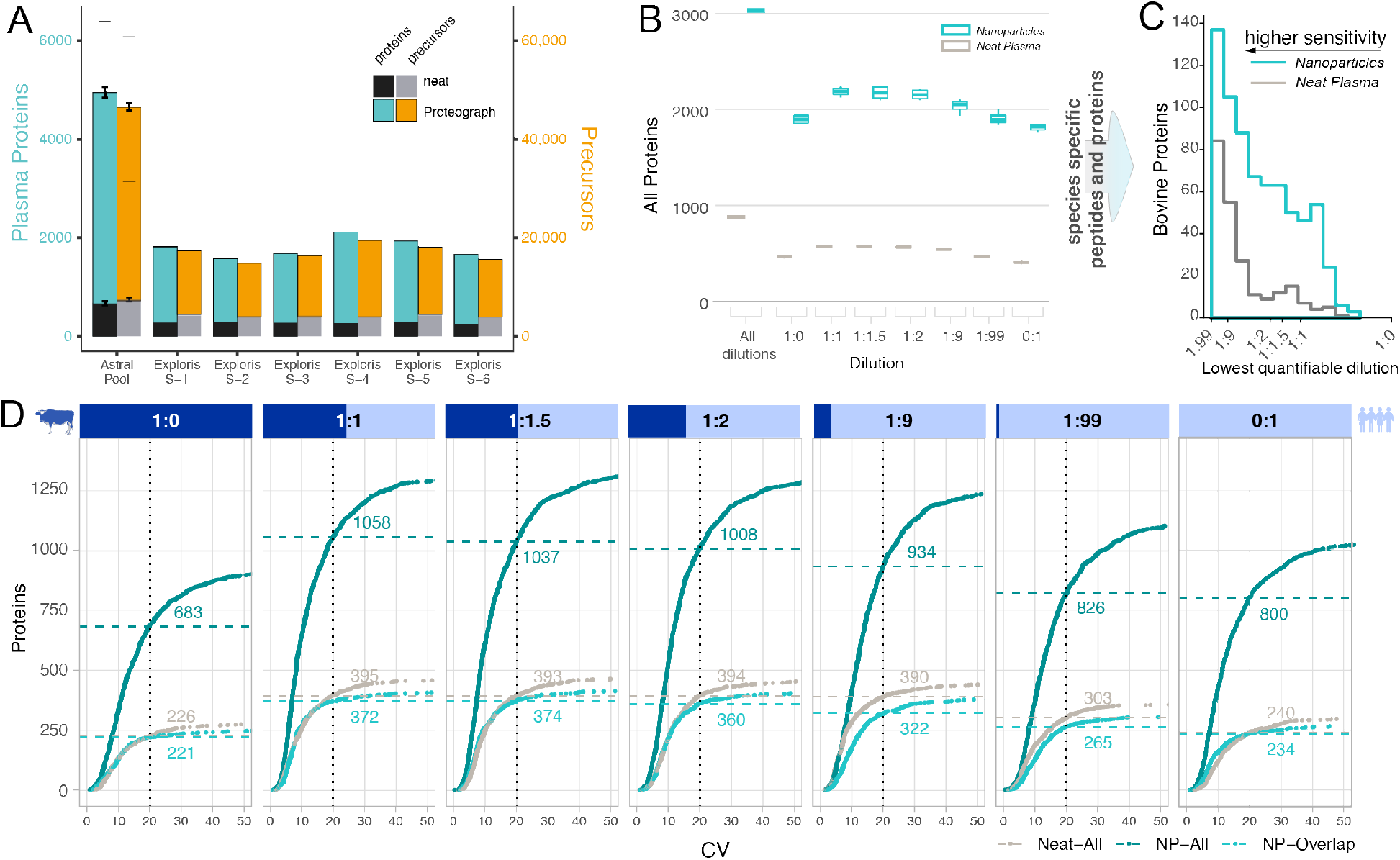
Identification and Precision Performance of a Neat Plasma and NP-corona Workflow. (A) Protein (left y-axis) and precursor identifications (right y-axis) determined for different LC-MS setups and biosamples comparing traditional neat workflows and NP workflow. Thermo Scientific Orbitrap Astral MS data was acquired for assay replicates standard deviations indicated by error bars, lower dash denoting identifications shared by all replicates (N=3), upper dash indicates identification across all replicates and nanoparticles. (B) The number of proteins quantified in mixed species plasma experiment on Exploris 480 at each ratio for the NP (teal) and neat workflow (grey), including bovine proteins and human proteins and those that are shared between species. Subsequent plots focused on species specific (unique) peptides and proteins. (C) Lower limits of quantification (LoQ) for bovine proteins quantified in both workflows. LoQ is the lowest level of dilution with a stringent quantitative response (lower is better). (D Number of proteins identified at a given coefficient of variation (CV) threshold for each spiked-in ratio. X-axis is the CV, and the Y-axis is the total number of bovine and human proteins quantified in four replicates with a CV lower than the given threshold. NP-workflow proteins also identified in neat (light teal), NP-workflow with all proteins (dark teal), and neat workflow (grey). Data shown is for IP10 human plasma pool.

### Fold-change Accuracy of Workflows

For accuracy and linearity evaluation we restricted our analysis to bovine-specific peptides and proteins as wider range of bovine concentrations measured allowed more robust estimations. To evaluate the relative fold-change accuracy of the NP workflow, we compared the measured fold-changes (observed change in protein intensities between spike-in ratios) with expected fold-changes (expected change in protein concentrations between spike-in ratios) of bovine proteins in 15 pairwise comparisons with 6 different spiked-in ratios. **Figure 4A, B** shows the results for three pairs of bovine spike-in ratios (2:1, 5:1, and 10:1), and the results for all 15 comparisons are shown in **Supplementary Figure 1**. The median fold-change is consistent across dilutions for both, the reference NP and neat digestion workflows, with a wider spread of fold-changes for the NP workflow. The expected fold-change is within the interquartile range (IQR) in most cases. For high dilutions, fold-change accuracy is increased for proteins quantified based on more than 1 peptide (**Supplementary Figure 1**) demonstrating the utility of quantifying multiple peptides per protein. Importantly, at any fold-change accuracy threshold NPs quantify more proteins than the reference neat plasma workflow (**Figure 4C, Supplementary Figure 2**). For the biosample PC6, we observed slightly decreased accuracy (**Supplementary Figure 3**). This pooled human blood plasma was collected differently than bovine plasma (CPD *vs*. K_2_EDTA) resulting in a variable buffer background across spike-in ratios. This appears to reduce accuracy and, since buffers mixing does not take place in clinical studies, we focus on IP10 (K_2_EDTA) for the remaining analyses.

**Figure 4.**
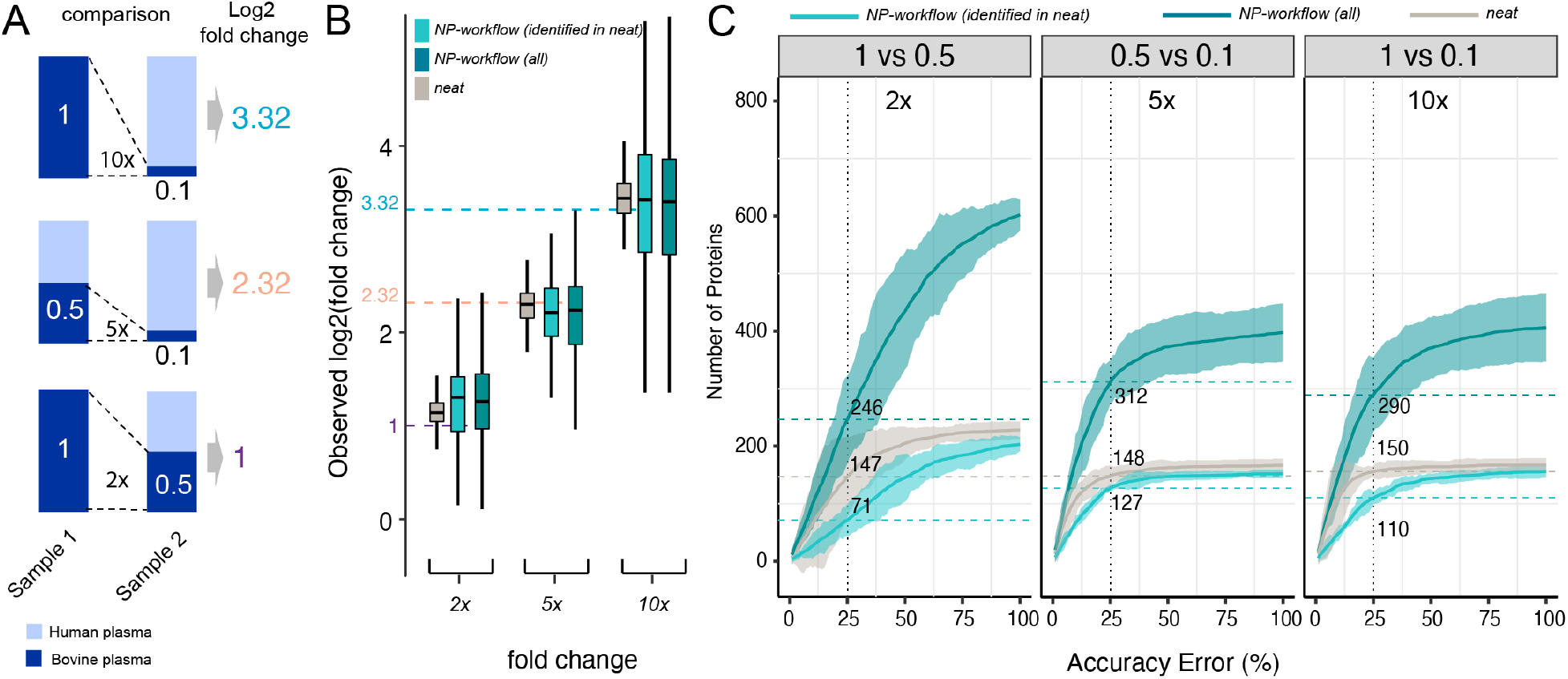
Fold-change Accuracy Performance of a Neat Plasma and NP-corona Workflow. (A) Three representative pairs of spiked-in samples and the expected fold-changes of bovine proteins concentration in these pairs. (B) Distribution of observed fold-changes of bovine proteins for 3 selected comparisons of spiked-in samples. The color indicates the data source: neat digestion (grey), Proteograph workflow (dark teal), or Proteograph workflow, but constrained to proteins also identified in neat (light teal). The horizontal dashed lines indicate the expected fold-changes. Boxplots report the 25% (lower hinge), 50%, and 75% quantiles (upper hinge). Whiskers indicate observations equal to or outside hinge ± 1.5 * interquartile range (IQR). Outliers (beyond 1.5 * IQR) are not plotted. (C) The number of bovine proteins identified at a given accuracy threshold for each expected fold-change. X-axis is the % accuracy error, i.e.,|log_2_ *FC*_*observed*_ − log_2_ *FC*_*expected*_|/log_2_ *FC*_*expected*_. The Y-axis is the number of proteins with an accuracy error below the given threshold. The horizontal dashed lines indicate proteins reported at the 25% threshold. Ribbon denotes the 99^th^ confidence interval. Data shown here are based on IP10.

A recently published study (15) compared a multi-step depletion workflow with neat plasma. Despite differences in biosamples, LC-MS methods, spike-in ratios, and computational pipelines, the fold-change distributions are comparable for similar spike-in ratios (**Supplementary Figure 4)**. Together, these results demonstrate NP-coronas capture quantitative differences across a wide range of concentrations and enable identification of substantially more proteins with similarly high quantification accuracy compared to a traditional neat plasma digestion workflow.

### Linearity of Quantitative Differences

Proteins that exhibit lower fold-change accuracy can still be highly valuable if they follow a systematic, e.g., log linear range trend, of quantification. The accuracy analysis (**Figure 4)** shows that, in comparison to a neat workflow, log fold-changes for a subset of proteins measured by the NP workflow can be compressed or inflated, due to the dynamic range compression that achieves much improved access to proteins in complex biosamples. We calculated the Pearson correlation between observed and expected log fold-changes for all proteins detected in the respective assay to evaluate linearity of the response, i.e., estimate how well protein regulation across biosamples can be captured. Across all spiked-in ratios (15 comparisons), we observed high linearity of the NP-workflow with a median Pearson correlation 0.995 that compares 0.998 in neat (**Figure 5A**). As well, most of the shared proteins, show a similar high linearity with slight advantages for the high abundance proteins detected in neat (**Figure 5A**).

**Figure 5.**
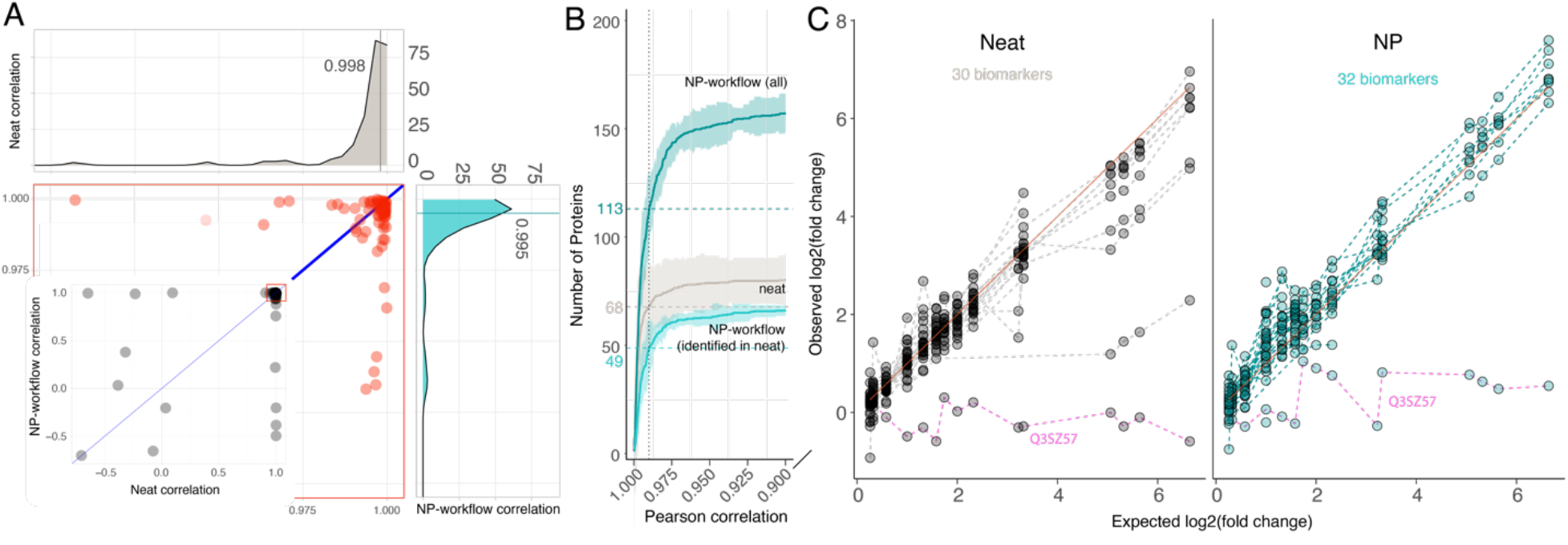
Linearity of Protein Quantification for Neat and Proteograph Workflows. Pearson correlation is calculated between observed and expected fold-changes of bovine proteins. (A) Neat digestion correlation versus Proteograph workflow correlation. Each dot represents one bovine protein. The marginal density plots show the distribution of Pearson correlation. 84 bovine proteins detected in all seven spiked-in ratios by both neat digestion (Grey color) and Proteograph workflows (Teal color) were plotted. (B) The number of bovine proteins identified at a given correlation threshold. X-axis is the Pearson correlation (truncated at 0.95), and the Y-axis is the number of bovine proteins with a correlation higher than the given threshold. The horizontal dashed lines indicate the number of proteins with a correlation ≥ 0.99. Proteograph workflow with proteins identified in neat workflow is colored in light teal, Proteograph workflow with all proteins is colored in dark teal, and neat digestion workflow is colored in grey. Data shown here are based on IP10. Plots depict bovine proteins only (identified with at least one bovine-specific peptide) that are detected at least once across all 15 pairwise comparisons. (C) Linearity of biomarkers matched to bovine proteins based on their gene symbols. 32 biomarkers were detected by the NP workflow while 30 biomarkers were detected in neat. Dashed lines connect the estimated fold-changes for each biomarker. Pink dashed line shows common outlier Q3SZ57. Selection of the most linearly responding peptides is based on Pearson correlation p-value determined for two assay replicates for matched biomarkers in neat workflow and NP-workflow. Depicted is the average of the two remaining replicates for the selected peptides.

In line with our observation for fold-change accuracy and consistent with an improved limit of quantification, the NP workflow quantifies more proteins at any linearity threshold. For instance, a stringent Pearson correlation of ≥ 0.99 was calculated for consistently detected, bovine-unique proteins across all dilutions. The NP workflow identified 113 such proteins, compared to 68 proteins detected by the neat plasma workflow (**Figure 4B**). Proteins demonstrating linearity with a Pearson correlation < 0.9 were minimal, 3 out of 83 for neat plasma, 25 out of 182 for NP, and 4 out of 70 for the shared proteins.

A key advantage of MS-based proteomics is to detect and quantify proteins through independent measurements of multiple peptides comprising unique amino acid sequences. This opens the opportunity to target peptides that present high linearity for efficient multiple reaction monitoring (MRM) MS acquisition. To compare quantitative performance of the ‘best’ peptides for human biomarker proteins mapped to bovine protein gene names, we selected the most linear peptide from two assay replicates based on Pearson correlation p-value. Subsequently, we evaluated linearity for these peptides in the remaining replicates (**Figure 5C**). The results demonstrated good linearity for both workflows, excluding Q3SZ57 (AFP), identified as a common outlier.

Despite the NP workflow’s design intent to significantly lower the signal (i.e., mass) for high-abundant proteins and thus, enhance the detection of low abundant content of proteomes, the signal maintained robust linearity for both proteins and peptides. This finding underscores NP’s utility for capturing proteins and quantifying dynamics across low and high abundance proteins (**Figure 4C, 5B,C**). Furthermore, the high degree of linearity indicates that further accuracy enhancements may be attained through additional normalization in both the neat plasma and NP-based workflows. Indeed, employing a simple linear fit correction for individual precursors, we were able to increase the fold-change accuracy in the NP workflow by a median of 34.1% across the dynamic range (**Supplementary Figure 5**).

### Statistical Power in Large-scale Studies

All steps of a proteomics workflow, including sample preparation, chromatography, and MS, can contribute to variation in results. To dissect individual steps in the workflow, we evaluated how CVs change with each workflow component (**Figure 6A**). For LC-MS reinjection alone, we observed a CV of 8.9% while the CV for the full workflow was 16.8% within batch and 20.9% across batches (including corona formation, protein digestion, peptide cleanup, different SP100 instruments and Proteograph Assay Kits on different days). Thus LC-MS accounts for about half of the overall variation (**Figure 6B**).

**Figure 6.**
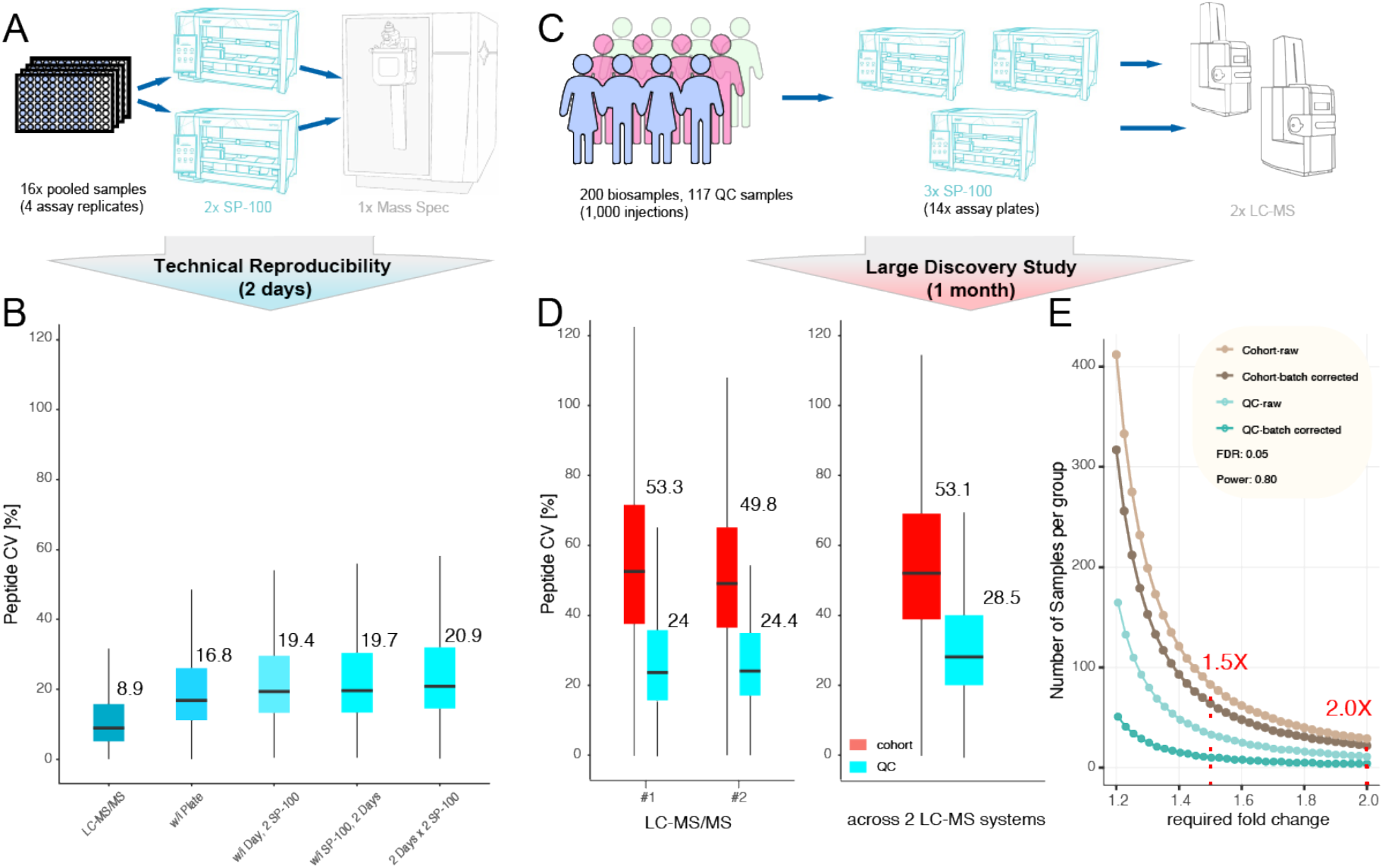
Reproducibility and Statistical Power for Proteograph Workflow in large-scale Discovery Proteomics Studies. (A) Experimental design of the reproducibility study. (B) Peptide level CV within and across plates, Proteograph SP100 automation instruments, and days of LC-MS analysis. (C) Experimental design of cohort study. (D) Peptide CV within and across LC-MS/MS instruments. (E) Statistical power analysis for a 200-sample, 1,000 injections cohort study detecting fold-change difference with 5% FDR and a power of 0.8.

To estimate the precision of NP-workflow across an entire biomarker study, we compared the performance for 117 plasma quality control (QC) samples versus 1,000 injections of 200 unique biosamples. This allowed us to compare QC-samples across 14 Proteograph assay (sample preparation) runs and peptides analyzed on two distinct LC-MS systems (**Figure 6C**). The QC variance accumulated over time to around 24% for a single LC-MS system and 28.5% across two LC-MS systems (**Figure 6D**). Importantly, this is substantially lower than the combination of biological and technical variation observed here and in other proteomics studies which is larger than 40% (41,42). We also estimated the statistical power to detect protein abundance differences based on the average variance across QC samples (28.5%), as well as patient samples (53.1%) with and without controlling for known assay variables such as sample preparation batch and LC-MS system. To detect biological differences of 2× at a stringent false discovery rate of 5% less than 30 samples (per group) are required in this cohort. If the biological background heterogeneity is low (e.g., for cell lines) the statistical power can depend more on technical variance. We approximated such cases considering the variance across all QC samples and determined that for detecting a 2× difference only 4 (controlling for assay variables) samples would be required, and 10 samples provide sufficient statistical power to detect fold-changes as little as 1.5× with high statistical confidence (**Figure 6E**). Together, this demonstrates sufficient reproducibility of the NP-workflow in large and small-scale studies to detect even small protein abundance differences.

## Discussion

Deep, unbiased access to the entire dynamic range of the plasma proteome at scale has great potential to advance our understanding of the proteome dynamics in health and disease. Previously, we have shown that NP-coronas compress the wide dynamic range of the plasma proteome, enabling deep and broad access to plasma proteins (3,14) at a scale to detect novel biomarkers (2,5). We have also shown that the high-throughput NP-based workflow provides quantitative response similar to ELISA assay, exemplified by four proteins (CRP, S100a8, S100a9, ANG) spiked in plasma at five different biologically relevant concentrations (2). Here, we conducted a comprehensive, proteome-wide evaluation of precision of quantification, fold-change accuracy, and linearity of log fold-changes as well as statistical power in cohort studies.

The utilization of multi-species plasma mixtures and the use of MS to evaluate quantification has several advantages over other approaches used to assess quantification and linearity. Firstly, measurements with ELISA may be subject to proteoform effects that are hard to distinguish from measurement error. For that reason, MS is used as a gold standard validation strategy for targeted affinity based capture assays (16,17). While NPs may sense proteoforms, the MS readout is more likely to resolve signal from multiple protein variants. Secondly, the multi-species approach allows us to simultaneously evaluate the quantification across many peptides of many proteins across the entire detection range rather than be limited to a few spiked-in proteins. Together, it allows us to more accurately estimate real world quantification accuracy in complex, plasma samples than would be achieved through individual measurements of isolated analytes.

The advantage of NPs compressing the dynamic range—i.e., transforming the absolute protein quantification (concentration)—is that it allows for unprecedentedly deep access to the dynamic range at scale without the need to target specific proteins of interest (unbiased approach). By design, NPs reduce the signal of high abundance proteins, such as albumin (signal compression), to capture more low abundance ones, like chemokines (signal enhancement). Our data demonstrate that, protein coronas not only render proteome content more visible to downstream detectors but also capture quantitative differences across samples with greater numbers of proteins and peptides at high precision, fold-change accuracy, and linearity. Moreover, the high linearity of the log fold-change response across a wide range of spiked-in ratios offers a clear path to combine NPs with rapid a MRM-based acquisition targeting highly quantitative peptides and further enhance fold-change accuracy by calibrating measured proteins and peptide intensities.

The feasibility of comprehensive biomarker discovery studies, spanning hundreds to thousands of plasma samples from human subjects or animal models, paves the way for the identification of a multitude of novel biomarkers. As we and other demonstrated previously, this is especially true within the traditionally challenging low-abundance range of the plasma proteome (4,18,19). Across such studies, high precision of quantification must be maintained, which we demonstrate in a 200-subject, 1000-sample pilot study is now possible using the automated NP workflow. Importantly, our analysis demonstrates that the technical variance even in large-scale studies comparing data across multiple instruments is substantially lower than the biological signal resulting in sufficient statistical power to discern even small biologically relevant protein abundance differences. Notably, not all biological variance is related to the phenotype under investigation and potential confounding factors may not always be identifiable or controllable (e.g., using models that control for age or BMI). This increases the necessity for larger-scale studies for complex human biology, underscoring the importance of workflows capable of accommodating thousands of samples. Collectively, our findings underscore the quantitative performance of protein corona-based assays, which holds significant opportunities for large scale, quantitative proteomics studies and biomarker discovery.

## Abbreviations

(NP): nanoparticle
(MS): mass spectrometry
(LC-MS): liquid chromatography mass spectrometry
(DIA): data-independent acquisition
(K_2_EDTA): etheylenediaminetetraacetic acid
(CPD): Citrate Phosphate Dextrose
(FA): formic acid
(ACN): acetonitrile
(LoD): Limits of detection
(CV): coefficient of variation
(IQR): interquartile range
(MRM): multiple reaction monitoring
(QC): quality control

## Authors Disclosure or Potential Conflict of Interest

O.C.F. has financial interest in Selecta Biosciences, Tarveda Therapeutics, and Seer where he is officer/director; and he serves as Senior Lecturer at BWH/HMS. S.F., Al.St., M.H., B.T., T.R.B., T.W., E.M.E., X.Z., E.S.O., A.A., B.L., J.C., M.F., J.W., M.G., H.X., C.S., Y.H., S.B., A.S., V.F., O.C.F., D.H. have financial interest in Seer, S.F., B.T., T.R.B., T.W., E.M.E., E.S.O., X.Z., T.W., J.C., M.F., J.W., M.G., H.X., C.S., A.S., V.F., O.C.F., D.H. have financial interest in PrognomiQ. E.D., T.A., A.H. are employed by Thermo Fisher Scientific. R.W. is a consultant to ModeRNA, Lumicell, Seer, Earli, and Accure Health. All other authors declare no conflicts of interest.

## Acknowledgment

The authors thank all members of the Seer Inc. team and Jinjun Shi for contributing critical discussions reviewing the manuscript. this work was supported in part by SBIR grant 5R44AG065051-02.

## Author Declaration

All authors confirmed they have contributed to the intellectual content of this paper with significant contributions to the conception and design, acquisition of data, or analysis and interpretation of data; drafting or revising the article for intellectual content; final approval of the published article; and agreement to be accountable for all aspects of the article thus ensuring that questions related to the accuracy or integrity of any part of the article are appropriately investigated and resolved.

