## Supplementary for "Protein Coronas on Functionalized Nanoparticles Enable Quantitative and Precise Large-Scale Deep Plasma Proteomics"

### Supplementary Methods

#### LC-MS Data Analysis Exploris 480 MS (supplementary information)

The main report with peptide quantifications was used for downstream analysis in R (v4.1). Raw precursor intensities were log-transformed and replicates of each spiked-in ratio were normalized using global median log-intensities normalization to correct for the technical variation among replicates. Ambiguous peptides shared by human and bovine proteins were removed from protein quantification to avoid mixing the signals from the two organisms. Log-transformed normalized precursor intensities were aggregated into protein group log-intensities using MaxLFQ (1) from R package "iq" v1.9.6 (2).

We first assessed the precision performance of different workflows by calculating the coefficient of variation (CV). That is, $CV = \frac{\sigma}{\mu}$ where $\sigma$ is the standard deviation, and $\mu$ is the mean of protein MaxLFQ intensities across the four replicates of each spike-in ratio.

We then evaluated the accuracy of the workflow by estimating how the measured protein fold changes between pairs of spiked-in ratios deviate from the expected ones. We focused on bovine proteins since their expected fold changes span from 1.2 to 100 (in comparison to 1.01–2 for human proteins). The spiked-in ratios for bovine proteins (1, 0.5, 0.4, 0.3333, 0.1, 0.01; samples with no bovine proteins were excluded) produce 15 pairwise comparisons with the following expected fold changes: 1/0.5 (2), 1/0.4 (2.5), 1/0.3333 (3), 1/0.1 (10), 1/0.01(100), 0.5/0.4 (1.25), 0.5/0.3333 (1.5), 0.5/0.1 (5), 0.5/0.01 (50), 0.4/0.3333 (1.2), 0.4/0.1 (4), 0.4/0.01 (40), 0.3333/0.1 (3.33), 0.3333/0.01 (33.33), and 0.1/0.01 (10).

Compression of dynamic range at the NP-protein interface can lead to systematic compression or inflation of the measured fold changes. To further improve quantification accuracy, one can correct these systematic deviations. We performed a linear fit between the expected and measured log fold-changes using the least square method for each precursor:

$$\log_2 \widetilde{FC}_i \sim \beta \log_2 FC_i + \varepsilon_i, \quad i = 1 \dots 15,$$

where $\log_2 FC_i$ and $\log_2 \widetilde{FC}_i$ represent the expected and measured $\log_2$ fold changes of the $i$-th comparison, respectively. The estimated linear fit parameter $\hat{\beta}$ was then used to calculate the corrected $\log_2$ fold change:

$$\log_2 \widehat{FC}_i = \frac{\log_2 \widetilde{FC}_i}{\hat{\beta}}$$

The errors between the expected and either measured or corrected fold-changes were calculated as $\frac{1}{15}\sum_i \left|\log_2 FC_i - \log_2 \widetilde{FC}_i\right|$ and $\frac{1}{15}\sum_i \left|\log_2 FC_i - \log_2 \widehat{FC}_i\right|$, respectively **(Supplementary Figure 5)**.

Finally, we evaluated the quality of protein intensities by matrix-matched calibration curve approach (3). For each analyte, we estimated the observed noise floor and linear intensity response to concentration by curve fitting. Limits of detection (LoD) and limits of quantification (LoQ) were estimated as the "spike-in ratio" (percentage of undiluted abundance) above which the predicted intensity response exceeds the observed noise floor by two standard deviations, and the concentration above which the coefficient of variation (CV) of intensity response (estimated by bootstrapping) falls below 20%, respectively. We required the background noise to be estimated from at least one concentration point (4 replicates), and the linear range to be estimated from at least two concentration points (8 replicates). To account for large/small steps at the extremes of our dilution range we employed a modified method that initially estimates noise from the lowest two concentrations and linear response by regression on the remaining points. This initial fit was refined by subsequent curve fitting and bootstrapping steps that matched the published implementation from (3).

**Cohort Study (supplementary information)**

The sample size that allows for detecting protein regulation at a given threshold ($\log_2 FC$) with given power (sensitivity) $\beta$, while also controlling the false discovery rate (FDR) (4,5) was calculated as:

$$n = \frac{2(z(1-{}^{\alpha}/_{2})+z(\beta))^2}{\left(\frac{\log_2 FC}{\sigma}\right)^2}, \text{ where } \alpha = \frac{\beta\, FDR}{1+\pi_0(1-FDR)},$$

$\sigma$ is the median standard deviation of $\log_2$ intensities for all proteins, $\pi_0$ is the ratio of non-regulated to regulated proteins in the data ($\pi_0 = 99$ was used), and $z(x)$ is the inverse cumulative density function of the standard normal distribution. We calculated the required sample size once without adjusting for known batch effects and setting $\sigma$ to the standard deviation of median normalized intensities for proteins present in 3 or more NP-specific samples, and once adjusting for known batch effects such as plate and LC-MS instrument using the residual variance from the following linear mixed model (in lme4 notation):

$$Abundance \sim 1 + (1|Plate) + (1|Instrument).$$

### Supplementary Figures

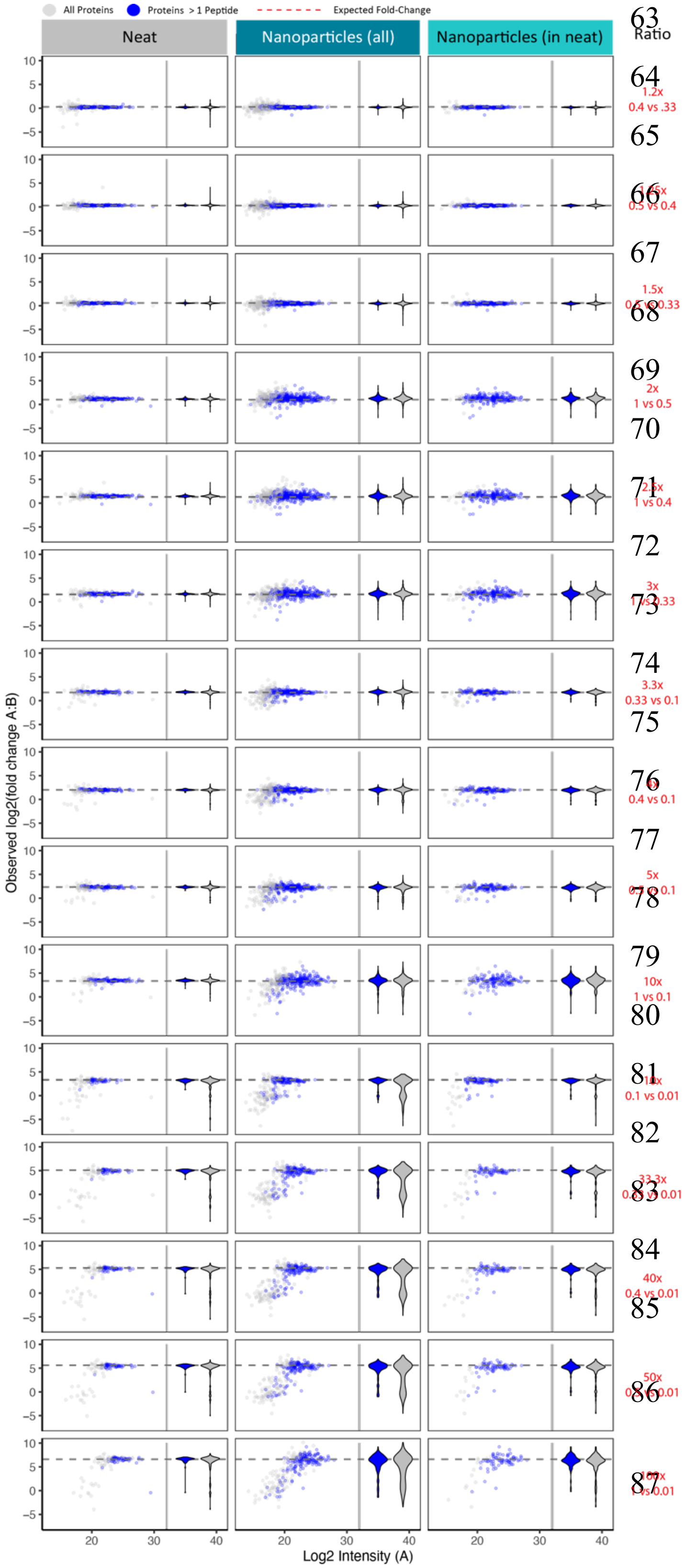


**Supplementary Figure 1. Protein Quantification Accuracy of Neat Plasma Digestion and Proteograph Workflows.** Spiked-in ratios for bovine proteins (1, 0.5, 0.4, 0.3333, 0.1, 0.01, and 0), producing 15 pairwise comparisons for small (A), medium (B), and large (C) fold changes. We skipped the ratio 0 since it had no bovine proteins. The pairwise comparisons were labeled as the expected fold-changes of the bovine proteins, i.e., 1 vs 0.5(2), 1 vs 0.4(2.5), 1 vs 0.3333(3), 1 vs 0.1(10), 1 vs 0.01(100), 0.5 vs 0.4(1.25), 0.5 vs 0.3333(1.5), 0.5 vs 0.1(5), 0.5 vs 0.01(50), 0.4 vs 0.3333(1.2), 0.4 vs 0.1(4), 0.4 vs 0.01(40), 0.3333 vs 0.1(3.33), 0.3333 vs 0.01(33.33), and 0.1 vs 0.01(10). X-axis denotes the log2 intensities. Y-axis is denoting fold changes of bovine proteins with dotted line indicating the expected fold change. For each comparison (neat, NP workflow all, shared), we illustrate all proteins as well (grey) as well as those quantified with more than 1 peptide (blue) as a scatter plot as well as violin plot. Data shown here are based on IP10.

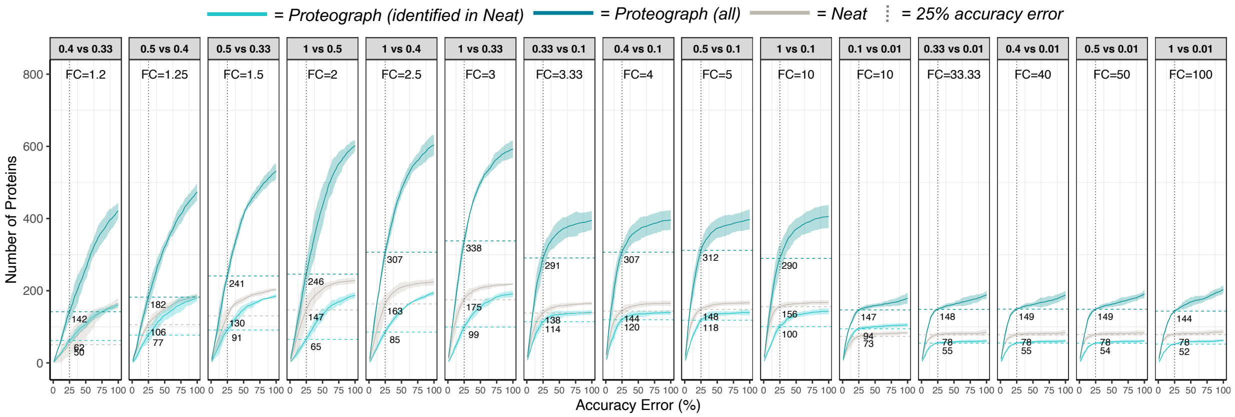


**Supplementary Figure 2. Quantitative Accuracy Performance of Proteograph Workflow in Comparison to Neat Digestion Workflow.** Each panel represents one fold-change of bovine proteins. X-axis is the % accuracy error, i.e., the difference between the observed and expected fold-change divided by the expected fold-change, and Y-axis is the number of bovine proteins identified at a given accuracy threshold. The horizontal dashed lines indicate bovine proteins reported at 25% threshold. Proteograph workflow with proteins identified in neat workflow is colored in light teal, Proteograph workflow with all proteins is colored in dark teal, and neat digestion workflow is colored in grey. Proteograph workflow with all proteins has demonstrated higher protein identification than neat digestion workflow with certain accuracy. Data shown here are based on IP10.

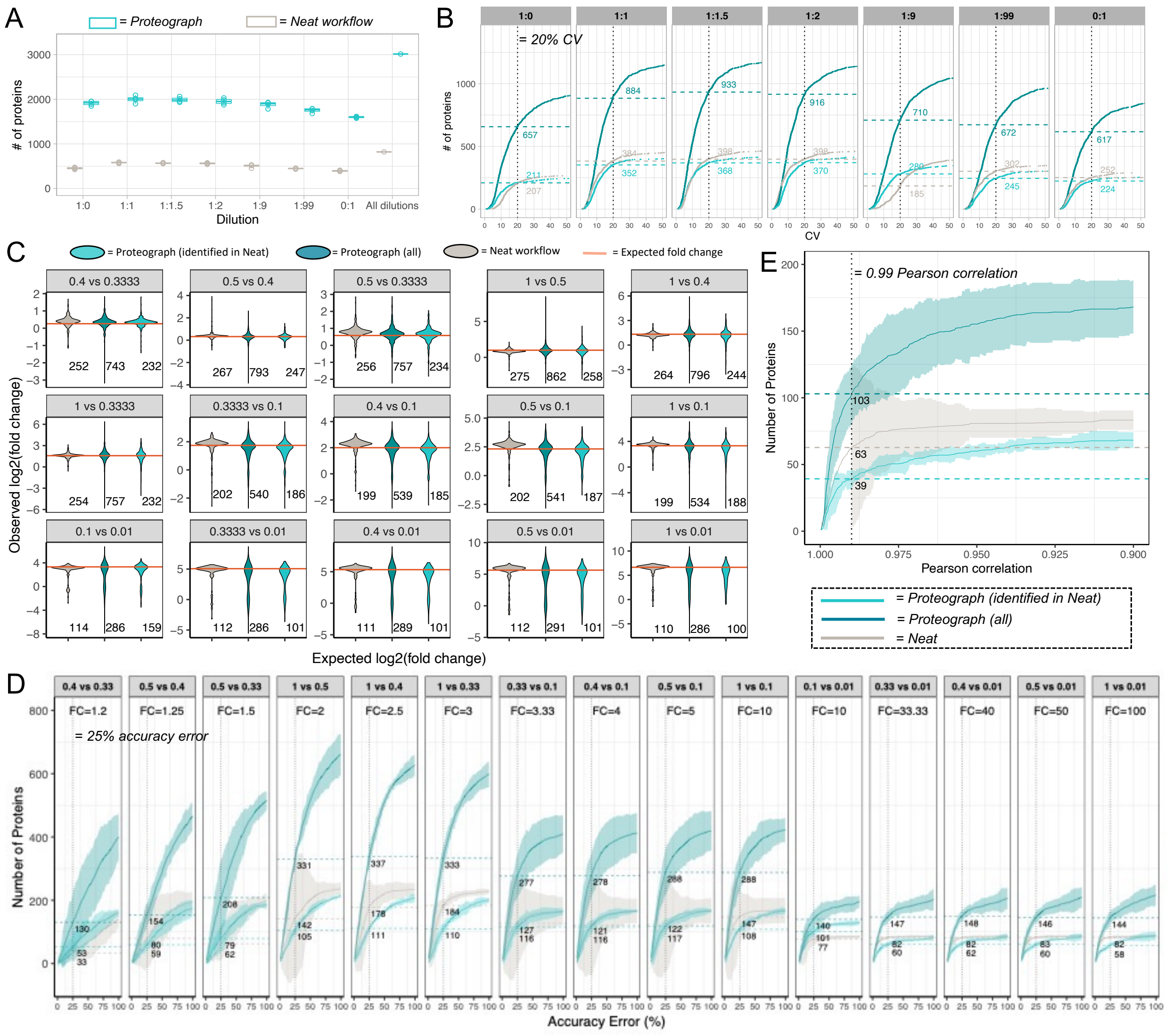


**Supplementary Figure 3. Quantification Performance of Neat and Proteograph Workflows for Another Human Plasma Sample PC6.** All the data processing and statistical analysis steps are the same as those on the previous IP10 sample. (A) The number of proteins quantified at each ratio. Proteograph workflow is colored in teal and neat workflow is colored in grey. (B) Number of Proteins Identified at a given CV Threshold for each Spiked-in ratio. X-axis is the CV calculated across four replicates, and the Y-axis is the number of proteins with a CV lower than the given threshold. NP-workflow with proteins identified in neat workflow is colored in light teal, Proteograph workflow with all proteins is colored in dark teal, and neat workflow is colored in grey. Only proteins quantified in all four replicates are counted. (C) Fold change accuracy of Neat and NP- Workflows. X-axis is the 15 comparisons and y-axis is observed fold changes. The orange dots connected by the orange line, as well as the orange numbers, indicate the expected fold changes of bovine proteins. The distribution of observed fold changes is shown by each box plot. The barplots at the top panel show the corresponding number of proteins summarized by each box. (D) Number of Proteins Identified at a given accuracy error for each expected fold change. Each panel represents one fold change. X-axis is the % accuracy error, i.e., the difference between the observed and expected protein fold change divided by the expected fold change, and Y-axis is the number of proteins identified at a given accuracy threshold. The horizontal dashed lines indicate proteins reported at 25% threshold. Proteograph workflow with all proteins has demonstrated higher protein identification than neat digestion workflow with certain accuracy.

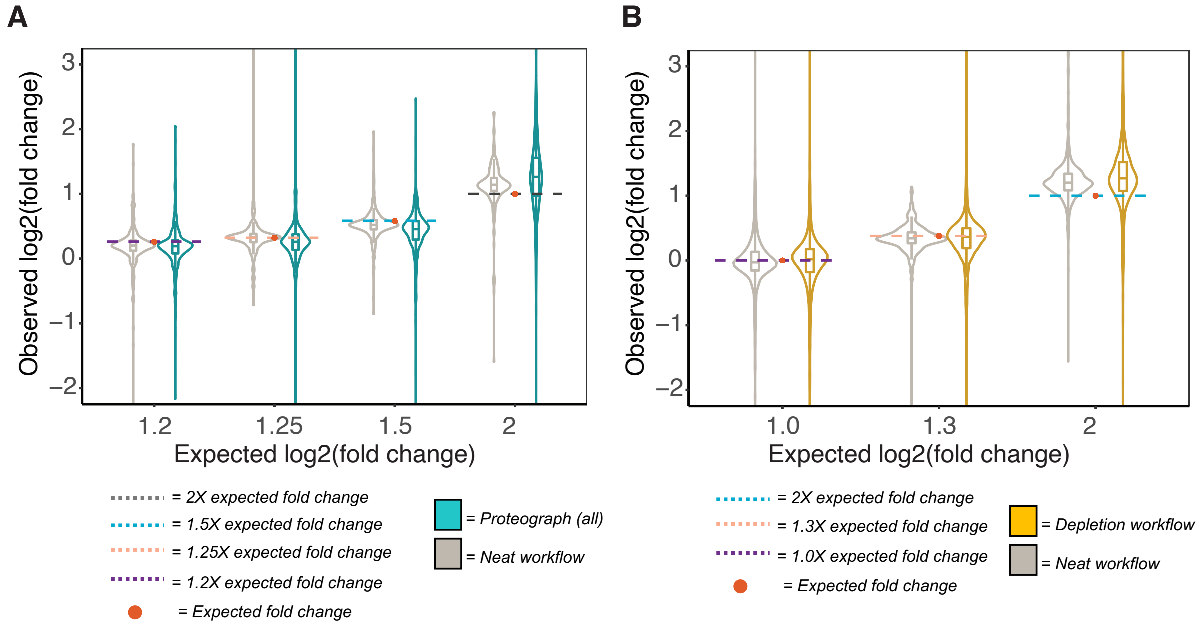


**Supplementary Figure 4. Quantification Accuracy Across Lower Fold Change Ranges.** (A) Proteograph workflow quantitation accuracy across 1.2-2 fold-change. Quantification data produced by NP-workflow shows comparable accuracy with those from neat workflow at low fold change of 1.2-2 fold. Data shown here are based on IP10. (B) The performance of depletion workflow is compared with neat digestion workflow here across similar low fold changes of 1.0-2 fold-change (6). Although this data is generated on different plasma sample and LC-MS workflow, overall we observe similar accuracies with Proteograph workflow.

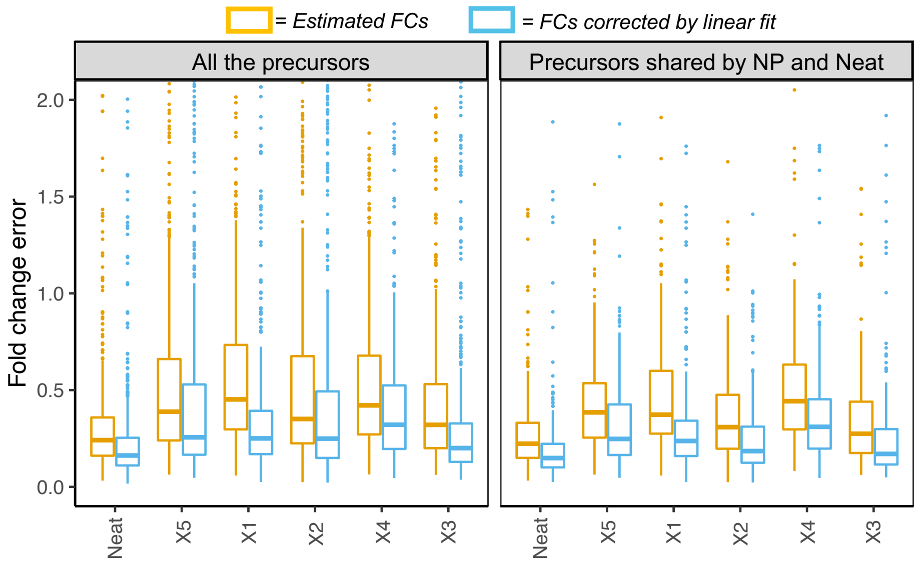


**Supplementary Figure 5. Quantification Accuracy Correcting for Systematic Shifts in Quantification.** For each precursor, we performed a linear fit correction between the measured log FC and the expected log FC of bovine precursors. The fold change error (y-axis) was then calculated as the mean absolute difference between the measured or corrected log FCs and the expected log FCs. The x-axis is the Neat workflow and the 5 NPs used by Proteograph workflow. After correction by linear fit, the fold change error can be significantly reduced.
